# Supervised Phenotype Discovery from Multimodal Brain Imaging

**DOI:** 10.1101/2021.09.03.458926

**Authors:** Weikang Gong, Song Bai, Ying-Qiu Zheng, Stephen M. Smith, Christian F. Beckmann

**Affiliations:** Centre for Functional MRI of the Brain (FMRIB), Nuffield Department of Clinical Neurosciences, Wellcome Centre for Integrative Neuroimaging, University of Oxford, Oxford, UK; Department of Engineering Science, University of Oxford, Oxford, UK; Radboud University Medical Centre, Department of Cognitive Neuroscience, Nijmegen, Netherlands; Donders Institute for Brain, Cognition and Behaviour, Radboud University Nijmegen, Nijmegen, Netherlands

**Keywords:** Multimodality, Brain imaging, UK Biobank, Imaging-derived phenotypes, Non-imaging derived phenotypes

## Abstract

Data-driven discovery of image-derived phenotypes (IDPs) from large-scale multimodal brain imaging data has enormous potential for neuroscientific and clinical research by linking IDPs to subjects’ demographic, behavioural, clinical and cognitive measures (i.e., non-imaging derived phenotypes or nIDPs). However, current approaches are primarily based on unsupervised approaches, without the use of information in nIDPs. In this paper, we proposed a semi-supervised, multimodal, and multi-task fusion approach, termed SuperBigFLICA, for IDP discovery, which simultaneously integrates information from multiple imaging modalities as well as multiple nIDPs. SuperBigFLICA is computationally efficient and largely avoids the need for parameter tuning. Using the UK Biobank brain imaging dataset with around 40,000 subjects and 47 modalities, along with more than 17,000 nIDPs, we showed that SuperBigFLICA enhances the prediction power of nIDPs, benchmarked against IDPs derived by conventional expert-knowledge and unsupervised-learning approaches (with average nIDP prediction accuracy improvements of up to 46%). It also enables the learning of generic imaging features that can predict new nIDPs. Further empirical analysis of the SuperBigFLICA algorithm demonstrates its robustness in different prediction tasks and the ability to derive biologically meaningful IDPs in predicting health outcomes and cognitive nIDPs, such as fluid intelligence and hypertension.

## I. Introduction

Large-scale population neuroimaging datasets, such as the data from UK Biobank, provide high-quality multimodal magnetic resonance imaging (MRI) data, with the potential for generating markers of psychiatric and neurodegenerative diseases and uncovering the neural basis of cognition through linking across imaging features to behavioural or genetic data [1]. However, such massive high-dimensional data make statistical modelling challenging due to their multimodal nature and cohort size. Therefore, instead of working directly from voxel-level spatial maps, it is becoming popular to reduce these maps into summary measures, sometimes referred to as “imaging-derived phenotypes (IDPs)” [2]–[4]. IDPs can be spatial summary statistics such as global and regionally averaged tissue volumes, while other IDPs can be measures of functional and structural connectivity or tissue biology. Building statistical models from an informative set of IDPs can significantly reduce the computational burden and, compared to working from voxel-wise data, has a similar or even improved signal-to-noise ratio for use in associations with non-imaging variables and predictive analysis linking to, e.g., behaviour and genetics [1], [2], [4].

Methods for deriving IDPs can be divided into two categories, one expert-knowledge-based and the other data-driven. The former approaches are typically concerned with extracting summary signals from pre-defined anatomy or functional brain atlases [5]. Although simple and efficient, this approach has a few limitations. First, the atlases may not be equally valid across different areas of the brain. For example, existing atlases typically provide fine-grained delineations across sensory cortices and less detailed across multimodal association cortices. These differences may result in increased inter-individual differences across different brain areas, potentially masking the signal of interest. Second, these regional characterisations are often derived from underlying features that may not appropriately map onto different data modalities. For example, atlases based on cytoarchitectonic features may differentially be suitable for IDPs reflecting regional cortical thickness but may be less suitable for summarising measures of functional connectivity. Furthermore, with multimodal data, expert-knowledge-based approaches typically ignore cross-modal relationships and thus have limited ability to capture continuous modes of variations shared by different modalities. Data-driven approaches for identifying IDPs, e.g., variants of unsupervised spatial dimensionality reduction techniques, may overcome the aforementioned limitations of expert-knowledge-based approaches. For example, independent component analysis (ICA) and dictionary learning (DicL) have been widely used to define “soft” brain parcellations in resting-state functional MRI analysis [6], [7]. They are based on arguably objective criteria such as maximising non-Gaussianity or minimising data reconstruction errors and can, in theory, be applied to a wide variety of different modalities. In a multimodal setting, FMRIB’s Linked ICA (FLICA) [8] is one approach for identifying continuous spatial modes of individual variations that are related to a range of behavioural phenotypes and diseases (e.g., lifespan development [9] and attention deficit hyperactivity disorder [10]). In our previous work, we developed BigFLICA, extending the original computationally expensive FLICA to handle much larger datasets such as UK Biobank [2]. These data-driven approaches have the advantages of being objective and considering cross-modal relationships, thereby revealing patterns that are ignored by expert-knowledge-based approaches [11], [12].

One of the primary applications of extracting imaging features as IDPs is predicting non-imaging derived phenotypes (nIDPs), including demographic, behavioural, clinical and cognitive measures from individuals. While the approaches listed above are designed to capture spatial modes of variation from the imaging data faithfully, they are not explicitly optimised for the latter prediction task. Incorporating the “target” nIDP information into IDP discovery, therefore, may benefit IDP extraction. Previous studies have proposed (semi-)supervised approaches for IDP discovery. For example, Qi et al. developed a multimodal fusion with reference (MCCAR) approach [13] and applied it to find multimodality modes related to schizophrenia [14] and major depressive disorder [15]. Zhou et al. also proposed a multi-modal latent space approach for early dementia diagnosis [16]. Another line of research focused on complex nonlinear approaches, such as multiple kernel learning [16]–[18], graph-based transductive learning [19] and neural networks such as multilayer perceptrons [20], [21], which proved successful in predicting neurological disorders such as Alzheimer’s disease. However, two caveats still exist in the above approaches. First, most of them do not scale well to big datasets due to expensive computational loads and high memory requirements. Second, nonlinear approaches heavily rely on parameter tuning and therefore require additional (cross-)validation for setting appropriate values. Furthermore, it is often difficult to make meaningful interpretations of the”black-box” nonlinear approaches, as explanations for deep neural networks produced by existing methods largely remain elusive and are yet to be standardised [22], [23]. As a result, it remains difficult to interpret the neural system that each feature (or spatial summary statistic) represents.

In this paper, to address these issues, we developed “Supervised BigFLICA” (SuperBigFLICA), a semi-supervised, multimodal, and multi-task fusion approach for IDP discovery, which simultaneously integrates information from multiple imaging modalities as well as from multiple nIDPs. By incorporating nIDPs in the modelling, one can hope to achieve better nIDP prediction than by training on the imaging data alone, as this exploits the covariance structure inherent in the nIDP space in addition to the predictive power of the imaging data. In the model, we use one or more target nIDPs to help the model learn spatial features that are biologically important in that they are generically useful in prediction, rather than only taking the route of classical unsupervised approaches of simply focusing on learning features for representing/reconstructing the image data with minimal loss. Further, using multiple nIDPs in training - a technique known as multi-task learning [24] - one can hope to refine the learned latent space better than when using single nIDPs, which are often noisy descriptions of the phenotype of interest (e.g., fluid intelligence). Compared with learning to predict each of the response variables individually, training across a range of noisy but related tasks simultaneously guides the model to characterise feature space shared across tasks, potentially leading to improved predictive power of the derived IDPs [24]–[26]. Additionally, the multi-task learning frameworks may still be useful even if one is interested in predicting unseen nIDPs (new tasks), because the latent space learned via a multi-task setting generically is more transferable and thus has higher predictive power.

SuperBigFLICA decomposes the imaging data into common “subject modes” across modalities, which characterise the inter-individual variation of a given underlying spatial component, along with modality-specific sparse spatial loadings and weightings [2], [8]. It minimises a composite loss function, consisting of both reconstruction errors of the imaging data (unsupervised learning) and the prediction errors of nIDPs (supervised learning), while additionally having constraints pushing for spatially sparse representations. Further, being built on a Bayesian framework, SuperBigFLICA can automatically balance the weights of different modalities and nIDPs, thereby aiming to largely bypass the need for parameter tuning. Optimised by a mini-batch stochastic gradient descent algorithm, SuperBigFLICA is computationally efficient and scalable to large datasets. In this study, we evaluate the performance of SuperBigFLICA across 39,770 UKB subjects, using 47 imaging modalities and 17,485 nIDPs. We provide a comprehensive empirical analysis of the SuperBigFLICA algorithm and demonstrate its potential for predicting health outcomes and cognitive nIDPs, and show that SuperBigFLICA enhances the predictive power of the derived IDPs and improves computational efficiency, benchmarked against IDPs created by conventional expert knowledge, or classical unsupervised or existing (semi-)supervised learning approaches.

## ll. Methods

### A. An overview of SuperBigFLICA

**Fig. 1** provides an overview of the SuperBigFLICA approach. SuperBigFLICA takes multimodal brain images and nIDPs as inputs, and learns a low-dimensional brain image derived feature set (data-driven IDPs) that is predictive of non-imaging derived phenotypes (nIDPs). This low-dimensional reduction is learnt as a linear combination of multimodal images, where the weights are sparse and represent the contribution of each voxel of each modality to each component. The low-dimensional feature set predicts multiple nIDPs using a sparse linear model. In other words, SuperBigFLICA uses linear layers for both image data reduction and nIDP prediction, with specific regularization designed for multimodal brain imaging data, to automatically balance the weights among modalities and nIDPs (tasks).

**Fig. 1:**
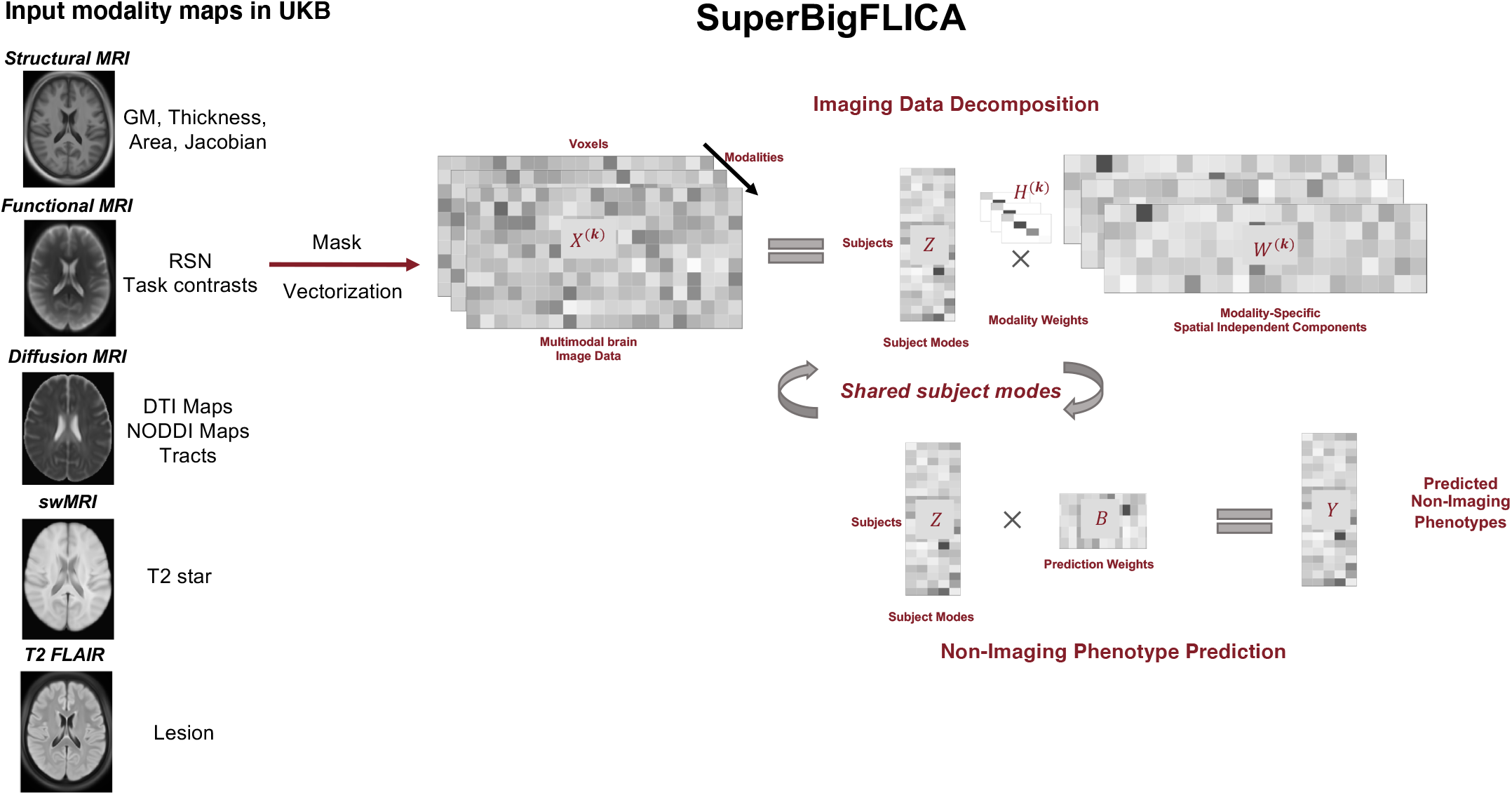
Overview of the workflow of the proposed SuperBigFLICA approach for (semi-)supervised multimodal fusion and phenotype discovery. From the left side, the inputs for multimodal analysis are 47 voxel-wise modality maps derived from 5 modalities in the UK Biobank dataset. These maps and nIDPs are then fed into the main SuperBigFLICA algorithm for multimodal learning. The final outputs are the predicted nIDPs. Abbreviations: GM: grey matter. RSN: resting-state network. DTI: diffusion tensor imaging. swMRI: Susceptibility weighted MRI.

### B. SuperBigFLICA from an optimisation perspective

We assume our data is being derived from a group of *N* subjects with multiple imaging modalities. Each of these modalities has been processed to produce one or more voxel-wise maps (or network matrices). For example, a task fMRI scan may produce several task contrast maps through statistical parametric mapping [27], and a diffusion MRI analysis can produce maps such as fractional anisotropy (FA) and mean diffusivity (MD) per subject. We assume that we have a total of *K* modality maps per subject, and each modality *k* is represented by a matrix **X**^(**k**)^ of size *N* × *P*_*k*_, where *P*_*k*_ is the number of feature values (e.g., voxels, tracts, areas, edges or vertices). We also assume that there are *Q* nIDPs per subject, summarised in a matrix **Y** of size *N* ×*Q*. We want to find an *L*-dimensional latent space across modalities, optimally predicting multiple nIDPs of interests in unseen subjects and representing the original imaging data. This latent space corresponds to the weights of continuous spatial modes representing inter-individual variations.

Formalising this in terms of a generative model, we will assume each modality map is generated as the product of the shared latent space, modality-specific spatial loadings and weights plus some Gaussian residual noise:

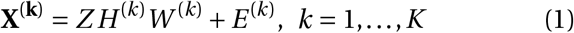

Meanwhile, the nIDPs of interest are generated by the product of shared latent space and the prediction weights plus some Gaussian residual noise:

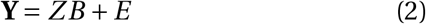

In the above two equations, 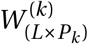 are the spatial loadings of the *k*-th modality map, which models the importance of each voxel to each latent dimension; 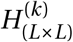 is a positive and diagonal modality weighting matrix (with 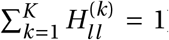), which reflects, for each component, the overall contribution of each modality; and *Z*_(*N*×*L*)_ is the latent features, i.e., the subject course shared across modalities; *B* is an *L*×*Q* matrix of prediction weights, which reflects the contribution of each latent dimension for predicting each nIDP; and finally, *E* ^(*k*)^ and *E* are independent Gaussian random error terms. **Fig. 1** shows an overview of the proposed approach.

We have three assumptions for the model. First, the subject loading *Z*_(*N*×*L*)_ is generated by:

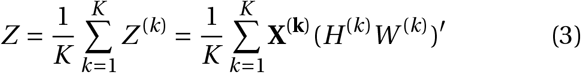

This shared subject loading *Z* is a weighted average of subject loadings per modality *Z* ^(*k*)^, in analogy to the original FLICA model [8]. This quantity is useful when we want to estimate the contribution of different modalities to the final prediction.

Second, the spatial loadings *W* ^(*k*)^ are approximately row-wise uncorrelated. We used a reconstruction loss to achieve this, which has been used in reconstruction independent component analysis [28]:

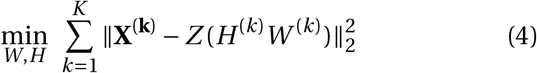

where *W* = (*W* ^(1)^,…, *W* ^(*K*)^), and *H* = (*H* ^(1)^,…, *H* ^(*K*)^). Note that the transpose of *H* ^(*k*)^*W* ^(*k*)^ in Eqn. (3) - due the soft orthogonality constraint for spatial loadings - will approximate the matrix inverse [28]. For example, when we have a single modality, the loss becomes 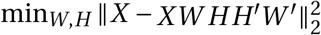, which means *WHH* ′*W* ′ will approximate the identity matrix. A similar property holds when we have *K* modalities.

Third, we assume sparsity in both spatial loadings and prediction weights - this we enforce through *L*_1_ regularisation. The orthogonal and sparsity constraints on spatial loadings will drive the model to find spatial sparse and non-Gaussian sources, similar to independent component analysis.

We combine the above model assumptions by means of the following objective function:

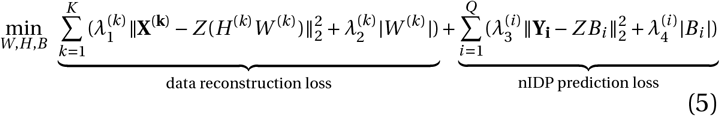

where *Y*_*i*_ is the *i* -th column of *Y*, and *B*_*i*_ is the *i* -th column of *B*.

### C. SuperBigFLICA in a Bayesian framework

Identification of the relative weighting parameters between the reconstruction loss, the prediction loss and the sparsity loss (i.e., the *λ* parameters in Eqn. (5)) through cross-validation is prohibitively expensive. Instead, we take a Bayesian perspective to tune the parameters. We assume normally distributed residual errors for the reconstruction and prediction terms and place Laplacian priors on the spatial loadings and prediction weights. Consequently, the *λ* parameters above will automatically become parameters in the distributions that can be jointly optimised with other model parameters.

For each imaging modality **X**^(**k**)^, the probabilistic model for the data reconstruction part is:

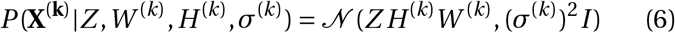

where *σ*^(*k*)^ is the modality-specific noise term (where we have assumed the noise is shared across voxels as in FLICA [8]). We place a Laplacian prior on each element of spatial loadings [29]:

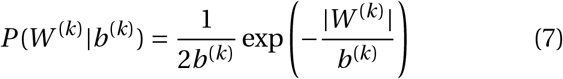

Further, we place a Gamma prior on each *x =* (*σ* ^(*k*)^)^2^ as 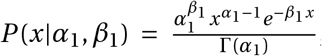, and a non-informative scale-invariant marginal prior on *x =* (*b*^(*k*)^)^2^ as *P* (*x*) *=* 1/*x*.

The probabilistic model for the prediction part is:

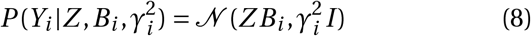

where *γ*_*i*_ is a task-specific noise term. We also place a Laplacian prior on prediction weights *B*_*i*_ [29]:

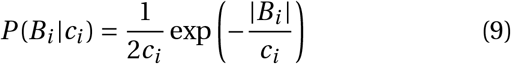

We place a Gamma prior on each *x =* (*γ* ^(*k*)^)^2^ as 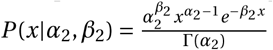, together with a non-informative scale-invariant marginal prior on 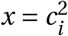 as *P*(*x*) = *1/x*.

The posterior distribution of model parameters *μ =* (*W* ^(*k*)^, *H* ^(*k*)^, *B*_*i*_, *σ* ^(*k*)^, *b*^(*k*)^, *γ*_*i*_, *c*_*i*_), *k =* 1,…, *K, i =* 1,…,*Q*, given the data *D =* (**X**^(**1**)^,…, **X**^(**K**)^, **Y**) then becomes:

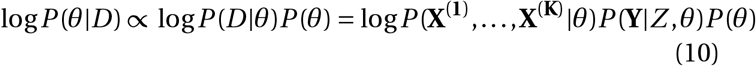

Note that the “auto” weights among imaging and nIDPs are proportional to the inverse of the residual prediction variance. In order to tune the model, we introduce a parameter *λ* ϵ [0, 1] to balance the weights between reconstruction and prediction losses. Tuning this *λ* is shown to be useful in different kinds of prediction tasks in our experiments later. Thus, we get a modified posterior to be maximized:

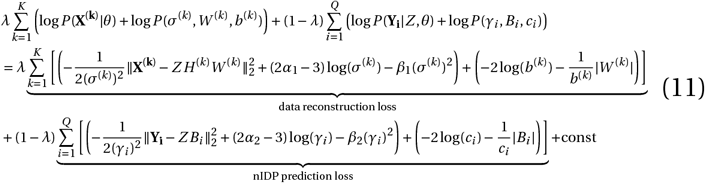

We can appreciate from Eqn. (11) that, the *λ*s in Eqn. (5) have been replaced by learnable parameters in the prior distribution of spatial weights and prediction weights (e.g., Gaussian and Laplacian priors), and these learnable parameters have their priors (i.e., Gamma or non-informative priors). The weights among modalities and nIDPs are proportional to the inverse of the residual prediction variance. This is analogous to a Bayesian linear regression with unknown residual variance [30]. The motivation is that the tasks (e.g., prediction or reconstruction) with larger error/uncertainty will be associated with lower confidence and therefore will be given lower weights [31]. We can further replace the Laplacian prior with other sparsity priors to achieve an equivalent sparsity effect. Such alternative priors include the automatic relevance determination (ARD) prior [32], the spike-and-slab prior [33], and the Gaussian mixture model prior (used in our original FLICA work [8]). We may later be interested in the contribution of a modality within a latent component to the prediction of a specific nIDP. This can be estimated by the correlation between a column of *Z* ^(*k*)^ in Eqn. (3) and an nIDP. This is because *Z* ^(*k*)^ is the subject course generated by the *k*-th modality, which has been used to predict an nIDP linearly.

### D. Model parameter optimization

The SuperBigFLICA model is implemented using Pytorch which can be easily run on a CPU and can be adapted to GPU usage for a more efficient model training. We obtain the maximum-a-posterior (MAP) solution of all parameters using a standard mini-batch stochastic gradient-descent (SGD) algorithms. Here, we have used the Adam optimizer [34] for parameters *W* ^(*k*)^, *H* ^(*k*)^, *B*_*i*_, and the RMSprop optimizer [35] for parameters *σ* ^(*k*)^, *b*^(*k*)^, *γ*_*i*_, *c*_*i*_, owing to empirical performance. The first order gradient-based algorithms were used because of their computational efficiency and low memory requirement suitable for optimising high-dimensional parameter space. Below, we evaluate the proposed combined optimisers with other standard first-order methods such as SGD with momentum [36], Adam or RMSprop, and a quasi-Newton methods L-BFGS [37]. We fixed the mini-batch size to 512 subjects and chose the optimal learning rate from 0.0001, 0.001, 0.01, and the tuning parameters *λ* from 0.99999 to 1*E* −5, in order to move from a pure data-driven to a purely supervised model. Dropout regularization with *p =* 0.2 is used on input modalities *X* ^(*k*)^ and subject loading *Z* to decrease the chance of overfitting [38]. Batch normalisation is used on *Z* in the training stage [39]. The total number of epochs (number of times the full data passes through the model) is 50, and the learning rate decreases by 1/2 every ten epochs. The model weights are initialised by Gaussian-distributed random numbers of mean 0 and variance 1.

### E. Connection and comparison with existing (semi-)supervised approaches

There exist some (semi-)supervised approaches that are somewhat similar to our SuperBigFLICA. The multi-site canonical correlation analysis with reference approach (MCCAR) [13] extends multi-site canonical correlation analysis to a supervised setting. MCCAR transforms each imaging modality into low-dimensional components, and simultaneously maximizes the correlation among components and the correlation between components and nIDPs. Using our notations, and re-defining *Z*^(*k*)^ as 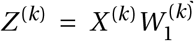, MCCAR’s objective function can be written as:

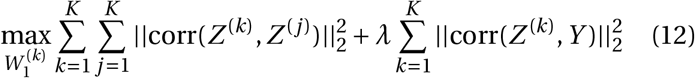

where corr(*A, B*) is the sum of column-wise correlation between matrix *A* and *B*. While MCCAR and SuperBigFLICA both aim to obtain subject loadings in a semi-supervised fashion, there remain three fundamental differences that may render MCCAR suboptimal for our purposes. First, MCCAR uses correlation to measure similarity. Therefore, it necessitates a full-batch iterative optimization approach for finding the best decomposition. Compared with Super-BigFLICA, the full-batch algorithm is not scalable to big datasets like UKB. Second, the number of terms in the objective functions scales quadratically with the number of modalities *K*, which further increases the computational complexity of optimization. Third, the weights among different modalities and nIDPs can not be determined automatically, and complex cross-validation is needed.

A more recent approach [16] (Zhou’s approach) uses a similar formulation as SuperBigFLICA. Using our notations above, the objective function of Zhou’s approach is:

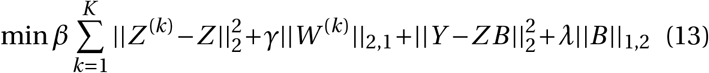

where ||*A*||_2,1_ and ||*A*||_1,2_ represent *l*_2,1_− and *l*_1,2_−norm of matrix *A* respectively. Zhou’s approach is similar to a non-Bayesian formulation of SuperBigFLICA, i.e., Eqn. (5), so that the weights among different modalities and nIDPs should be determined by cross-validation. In addition, Zhou’s approach uses the Augmented Lagrange Multiplier algorithm to optimize the objective function, which again requires a full-batch algorithm. Therefore, the scalability of Zhou’s approach to big datasets is limited. Similarly, Zhou et. al [40] also proposed another approach that uses an additional Laplacian regularizer to ensure that similar inputs have similar latent feature representations, which has similar disadvantages as the above-mentioned approach.

We compared SuperBigFLICA with two existing approaches mentioned in the Method section II-E. We compared the prediction correlation of 12 nIDPs using a 25-dimensional decomposition of different methods. We implemented the MCCAR approach [13] based on the FIT toolbox (https://github.com/trendscenter/fit). We implemented Zhou’s approach from scratch because the original code is not openly accessible [16]. We generated two variants of Zhou’s approach based on our SuperBigFLICA code. In the first version, we removed the auto-weighting parameters in SuperBigFLICA, to mimic the situation that Zhou’s approach is not able to automatically determine the weights between different modalities and tasks. In other words, the *æ*^(*k*)^, *b*^(*k*)^, *γ*_*i*_, *c*_*i*_ parameters in Su-perBigFLICA remain fixed, making the weights equal across modalities and tasks. In this case, we would assume that manually determining the optimal weighting parameters by cross-validation is computationally infeasible for datasets like UKB. In the second version, we further implemented the *l*_2,1_-norm regularization on spatial weights *W* ^(*k*)^. Note that in both implementations, SuperBigFLICA and Zhou’s approach have the same loss function on the data reconstruction and nIDP prediction parts.

For comparing the different methods, we choose a 25-dimensional decomposition because the MCCAR approach is computationally infeasible for higher dimensions in the UKB dataset in the authors’ implementation. Similarly, Zhou’s approach would theoretically suffer from the same computational issue. In our implementation of Zhou’s approach, we used SGD to address this problem.

## III. Experiments

### A. Brain imaging data

Voxel-wise neuroimaging data of 47 modalities from 39,770 subjects were used in this paper, including: (1) 25 modalities from the resting-state fMRI ICA dual-regression spatial maps after *Z*-score normalisation [1] ; (2) 6 modalities from the emotion task fMRI experiment: 3 contrast (shapes, faces, faces>shapes) of *Z*-statistics and 3 *contrasts of parameter estimate* maps [1] that reflect %BOLD signal change; (3) 10 diffusion MRI derived modalities (9 TBSS features, including FA, MD, MO, L1, L2, L3, OD, ICVF, ISOVF [41], [42] and a summed tractography map of 27 tracts from AutoPtx in FSL [43]); (4) 4 T1-MRI derived modalities (grey matter volume and Jacobian deformation map (which shows expansion/contraction generated by the nonlinear warp to standard space, and hence reflects local volume) in the volumetric space, and cortical area and thickness in the Freesurfer’s fsaverage surface space; (5) 1 susceptibility-weighted MRI map (T2-star); (6) 1 T2-FLAIR MRI derived modality (white matter hyperintensity map estimated by BIANCA [44]) (**Table I**). The UK Biobank imaging data were mainly preprocessed by FSL [45], [46] and FreeSurfer [47] following an optimized processing pipeline [48] (https://www.fmrib.ox.ac.uk/ukbiobank/). From the voxel-wise modality maps, our team generates 3,913 “expert-designed” (i.e., not data-driven) IDPs (which are disseminated by UK Biobank), covering the entire brain and including multimodal information on regional and tissue volume, cortical area, cortical thickness, regional and tissue intensity, cortical grey-white contrast, white matter hyperintensity volume, regional T2 star, WM tract FA, WM tract MO, WM tract diffusivity, WM tract ICVF, WM tract OD, WM tract ISOVF, tfMRI activation, rfMRI node amplitude, and rfMRI connectivity, from the same six imaging modalities we used in SuperBigFLICA (T1, T2-FLAIR, swMRI, task fMRI, resting-state fMRI, and diffusion MRI). The details of the IDP generation are illustrated in our previous work [1], [4].

**TABLE I:**
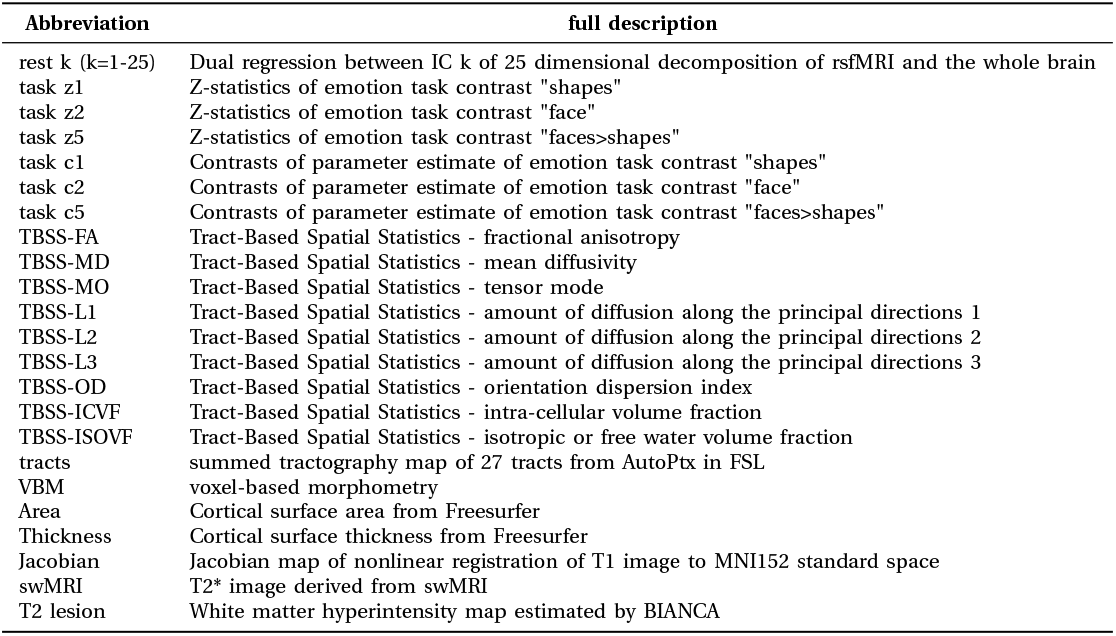
A description of 47 modalities of UKB dataset used in this paper.

### B. Non-imaging derived phenotypes

A total of 17,485 non-imaging derived phenotypes (nIDPs) were used in this paper. In the prediction tasks, we mainly analyzed 12 of them in the “physical”, “cognition”, and “health outcome” domains, summarised in **Table II**. These 12 nIDPs were selected to allow for a direct comparison to our previous work, as they were the best predicted nIDPs in cognition and health outcome domains by our baseline approach BigFLICA [2]. The direct comparison of performance between SuperBigFLICA and BigFLICA approaches enables us to study the benefits of including the supervised loss terms.

**TABLE II:** Non-imaging derived phenotypes used in this study.

| Non-imaging derived phenotypes |
| --- |
| Fluid intelligence |
| Hypertension |
| Handedness |
| Treatment/medication code (1140884600 - metformin) |
| Diabetes diagnosed by doctor |
| Non-cancer illness code, self-reported (1261 - multiple sclerosis) |
| Overall health rating |
| Age started wearing glasses or contact lenses |
| Number of treatments/medications taken |
| Mean time to correctly identify matches |
| Maximum digits remembered correctly |
| Number of self-reported non-cancer illnesses |

### C. Confounding variables and missing values

Before carrying out nIDP prediction, a total of 597 confounding variables were regressed out from both voxel-wise imaging data and nIDPs, using linear regression [49]. Missing modality data for a subject were imputed by the mean map of all other subjects. We did not impute missing nIDPs, and therefore, in the SuperBigFLICA model, only data-reconstruction-related losses play a role for subjects with missing nIDP data.

### D. Imaging data pre-reduction using dictionary learning

There are tens of thousands of voxels per modality, so a direct fitting of SuperBigFLICA using voxel-level data is computationally expensive and memory-consuming. Although we can perform mini-batch optimisation on the subjects, we need to keep *K* big voxel-by-component spatial maps as learnable parameters in memory. One possible solution is to use sparse dictionary learning for voxel space dimension reduction before running SuperBigFLICA. As shown in our previous work [2], for BigFLICA, sparse dictionary learning will reduce the computation load of the optimisation and may increase the modality-specific signal-to-noise ratio. This is because the overall model is linear, so a pre-dimension reduction using a (linear) dictionary learning should not interfere with important information that can be captured by BigFLICA, but will potentially have the de-noising effect. Owing to the similar property of the models employed in BigFLICA and SuperBigFLICA, we expect sparse dictionary learning to perform similarly well. Here, we evaluate the effect of data reduction on the final prediction across the voxel-domain between 100 and 2,000 dictionary features per modality before running any subsequent algorithm, e.g., BigFLICA or SuperBigFLICA. Note that this pre-reduction can also be performed with nIDP information included. We therefore also tested applying a “single-modality” SuperBigFLICA to each single modality map (which is just a special case with *K =* 1) with all 17,485 nIDPs as supervision before using SuperBigFLICA for multimodal analysis.

### E. Baseline: nIDP prediciton using hand-curated IDPs and modes of BigFLICA

In real data, we rely on the performance of predicting nIDPs as a surrogate criterion to evaluate different methods, given that “ground-truth” IDPs do not in general exist. As a baseline approach, we compare hand-curated IDPs and modes of BigFLICA. The pipeline for prediction follows our previous work [2]. In brief, 3,913 IDPs and 1,000 modes of BigFLICA are extracted from UK Biobank imaging data. BigFLICA used a 3,500-dimensional MIGP approach [2] and a 2,000-dimensional dictionary learning approach as data preprocessing [2] before running the core FLICA variational Bayesian optimization.

Here, a high dimensional decomposition was chosen as in our previous work on BigFLICA, which achieved the best prediction accuracy for most nIDPs [2]. Further, elasticnet regression, from the glmnet package [50], was used to predict each of the 12 nIDPs separately (known as single-task learning) using IDPs or FLICA subject modes as model regressors (features). This approach is widely used and has been shown to achieve robust and state-of-the-art performance in many neuroimaging studies [51], [52].

We randomly selected a subset of 25,000 subjects for model training, while the validation set contains 5,000 different subjects, and the test set was formed from the remaining 9,770 subjects. All methods’ comparisons are using the same train-test split. For single-task learning, the prediction accuracy was quantified as the Pearson correlation coefficient *r* between the predicted and the true values of each nIDP in the test sets. For multi-task learning, the prediction accuracy was quantified as the sum of correlations with nIDPs larger than 0.1. For comparing different methods, we used a bootstrap approach [53] to estimate the statistical significance of the difference of *r* between the two methods. In this approach, we resample the predicted nIDPs of the first approach in the test set with replacement, and recalculate the *r* -value of this approach 10,000 times, getting a bootstrap distribution of *r*. Then we estimate the proportion of observed *r* of the second approach that is larger than the bootstrap *r* -values. If it is larger than 95% of the bootstrap *r* -values, we will conclude that the second approach works significantly better than the first approach.

### F. Predicting nIDPs using single-task and multi-task SuperBigFLICA

To demonstrate how SuperBigFLICA with one or more target nIDPs in training can boost the performance compared to hand-curated IDPs and unsupervised BigFLICA, first, we trained single-task SuperBigFLICA with each of the 12 nIDPs as a supervision target. We then trained multi-task SuperBigFLICA by including each target and the top 24 or 99 most highly correlated (with the target) nIDPs from all 17,485 available nIDPs from the UK Biobank dataset, in the training stage. In previous work, it was already established that the inclusion of correlated tasks as targets could boost the performance of single-task learning [24], [25]. Therefore, we performed 25- or 100-dimensional multi-task learning (separately) for each of the 12 nIDPs and evaluated the prediction performance. Note that while an additional 24 (or 99) nIDPs (that are correlated to the target nIDP) are used to help in the training, they are not used in test data to help the prediction - only the imaging data from test subjects is used in predictions of nIDPs in test subjects. Finally, we trained SuperBigFLICA with all 17,485 nIDPs as supervision targets. For the tuning parameters, the number of latent components was chosen to be 25, 100, 250, 500, or 1,000, and the *λ* parameter is chosen from 1*E* − 5 to 0.99999. We evaluated the influence of *λ*, and different random model parameter initialisations on the final prediction performance.

Furthermore, we performed a more comprehensive comparison of prediction performance using IDPs, BigFLICA and single-task SuperBigFLICA on each of the 17,485 nIDPs. Owing to computational limitations, we fix some parameters of SuperBigFLICA to reduce the overall computation time, with a 25-dimensional decomposition and learning rate of 0.001, and the *λ =* 0.5 based on their empirical good performance in the 12 cognitive and health outcome nIDPs.

### G. Evaluation of the generalisability of SuperBigFLICA modes on unseen nIDPs

One of the fundamental goals of phenotype discovery is to learn a low-dimensional latent space that is generalisable in that it can predict “unseen” nIDPs. In the above analyses, the nIDP to be predicted was always included in the training stage. Here, we evaluated whether SuperBigFLICA can generate a good representation for entirely new tasks, wherein in the training stage, we only use nIDPs that are not our targets. To do this, for a given nIDP that we want to predict (e.g., fluid intelligence), we first compute the correlations between this target nIDP and all other 17,485 nIDPs. We selectively include the 16,485 least correlated nIDPs for training SuperBigFLICA, ignoring the 1,000 most strongly covarying nIDPs in training. This means that, for example, the mean correlation of 16,485 nIDPs with the target variable fluid intelligence is 0.007, and nIDPs with correlation *r >* 0.032 (*p <* 2 × 10^−7^,*n =* 25, 000) with fluid intelligence are removed in training. This experiment simulates a “bad” situation where almost no nIDPs are related to our target when generating the latent space. We then used elastic-net regression to predict our target nIDP using the resulting latent space, to evaluate the degree to which the inclusion of unrelated nIDPs constraints the learning towards IDP features that are more generally useful for prediction across a wide range of nIDPs, and thereby ultimately also improves prediction for specific nIDPs of interest. This may also relate to the concept of transfer learning, where we can use multiple nIDPs to learn a space that generically has high predictive power.

## IV. Results

### A. Comparing SuperBigFLICA with hand-curated IDPs and modes of unsupervised BigFLICA

We first compared SuperBigFLICA with hand-curated IDPs and BigFLICA, and then compared different variants of SuperBigFLICA, in terms of prediction accuracy of nIDPs (Experiments sections III-E, III-F and III-G). Each subfigure of **Fig.2** shows the overall prediction accuracy of different experimental approaches for 12 nIDPs. We report the results that use the dictionary dimension of 250 for each modality (detailed in the next section). The test accuracy is obtained using the best tuning parameters (*λ* and number of latent components) selected in the validation set.

**Fig. 2:**
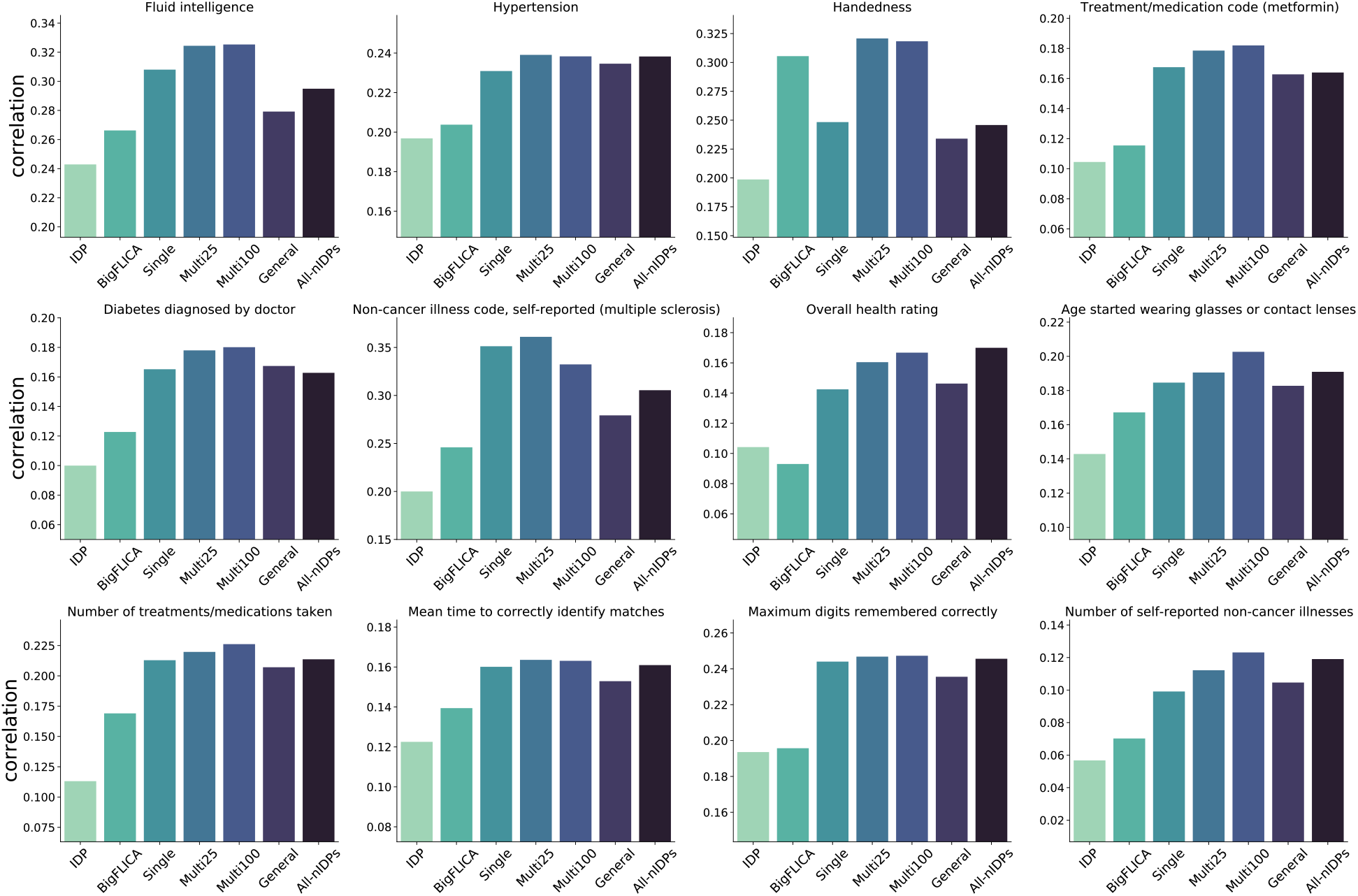
Comparison of SuperBigFLICA with hand-curated IDPs and modes of unsupervised BigFLICA for the 12 health outcome and cognitive nIDPs listed in Table II. Each figure shows - for a different nIDP - the predictive correlation of different approaches and different parameter settings. The first and second column shows the ‘baseline’ prediction accuracy of IDPs and BigFLICA. The third column shows the accuracy of single-task SuperBigFLICA trained end-to-end. The fourth and fifth columns show prediction accuracy of multi-task SuperBigFLICA, with 24 and 99 most correlated nIDPs as auxiliary tasks for supporting the training of the main nIDP of interest. The sixth column shows the generalisability accuracy of SuperBigFLICA modes on unseen nIDPs, where the main nIDP of interest is not included in the learning of latent space but only 16485 nIDPs that are least correlated with it. The seventh column shows the prediction accuracy when we fuse all 47 modalities and use all 17,485 nIDPs to train a single model.

The first two columns are the accuracy of elastic-net regression with hand-curated IDPs and modes of unsupervised BigFLICA as input features, while the third column is the accuracy of single-task SuperBigFLICA trained end-to-end. We can see that in almost all cases, the accuracy of (semi-)supervised training outperforms hand-curated IDPs and unsupervised BigFLICA. The average percent improvement of single-task SuperBigFLICA compared with hand-curated IDPs and BigFLICA are 46% and 25%. We further report the statistical difference of improvements. As shown in the first two columns of **Table. III**, the BigFLICA approach performs better than the IDP approach significantly, and the SuperBigFLICA approach also performs better than the BigFLICA approach significantly.

**TABLE III:**
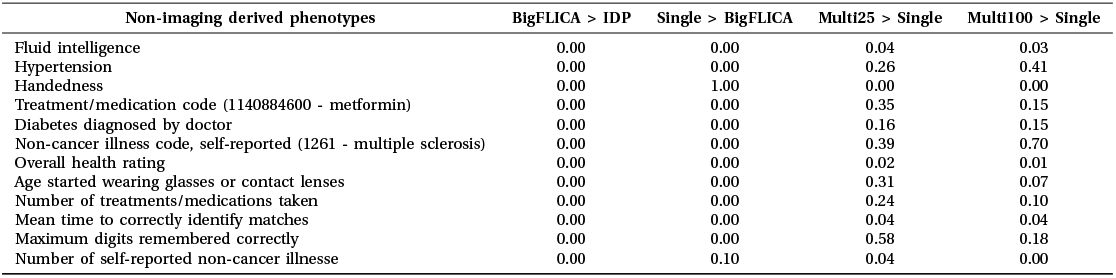
The p-values for comparing different methods.

The fourth and fifth columns of each subfigure of **Fig.2** show prediction accuracy of multi-task SuperBigFLICA, with 24 and 99 most correlated nIDPs as auxiliary tasks for supporting the training of the main nIDP of interests. We can see that multi-task learning usually further improves the prediction accuracy compared with single-task SuperBigFLICA. The average per cent improvement of two multi-task BigFLICA compared with single-task SuperBigFLICA are 9% and 7%. We further report the statistical difference of improvements. As shown in the last two columns of **Table. III**, multi-task SuperBigFLICA outperforms singletask SuperBigFLICA approach significantly in 5 of the 12 nIDPs (*p <* 0.05). As most health outcomes and cognitive nIDPs in our experiments are noisy, adding more noisy nIDPs to the joint multi-task training will only have a small benefit, thus, the improvement may not be statistically significant.

The sixth column of each subfigure of **Fig.2** shows the generalisability of SuperBigFLICA modes on unseen nIDPs, where the main nIDP of interests is not included in learning the latent space (only nIDPs that are at best weakly correlated with the main nIDP of interests are involved). We then used elastic-net regression to predict the main nIDP based on the learned latent space as regressors (We are still using the imaging data to perform nIDP predictions). Overall, this is expected to perform worse than single-task and multitask SuperBigFLICA because the target nIDP is not involved in the training. However, it still outperforms unsupervised BigFLICA plus elastic-net by 21%, and is slightly worse than single-task SuperBigFLICA. This experiment demonstrated that the inclusion of even irrelevant tasks in the “supervised” training could boost the predictive performance of the generated latent space.

The seventh column of each subfigure of **Fig.2** shows the prediction accuracy when we fuse all 47 modalities and 17,485 nIDPs to train one multi-task SuperBigFLICA model. We can see that the prediction accuracy is similar to singletask SuperBigFLICA for most of the nIDPs.

We tested the robustness of the above results. First, we tested the influence of adding uncorrelated nIDPs into multitask learning. We added the 25 most uncorrelated auxiliary nIDPs together with the 24 most correlated nIDPs for supporting the training of the main nIDP of interests. **Fig. 3** shows that prediction correlations remain virtually identical, except for the handedness nIDPs, suggesting the robustness of the model in accounting for noise in nIDPs. Second, we run the dictionary learning preprocessing only using the training set data (instead of on the whole imaging dataset), and to replace the fix train-test split with a threefold cross validation approach. **Fig. 4** shows the prediction correlation of 12 nIDPs using single-task and multi-task SuperBigFLICA, with and without the new settings. Here, the new correlations are computed as a mean of 3 folds. We can see that, except for one nIDPs, Non-cancer illness code, there are no statistical differences between the original settings and the new settings. We believe the results are robust owing to the large sample size we used; dictionary learning is not over-fitting, even if derived from test data. Third, we calculated the prediction performance of 12 nIDPs with missing values directly removed from the training process. In the original setting, we kept missing values in nIDPs or imaging data in the model fitting process, because the loss function has both reconstruction and prediction terms, we would expect, e.g., a subject with miss nIDP data still can contribute to the minimization of image reconstruction term. In the current setting, we just remove all subjects with either nIDP or imaging data missing. **Fig. 5** shows that compared with including the missing values in the training, removing them directly in the training has a very similar performance.

**Fig. 3:**
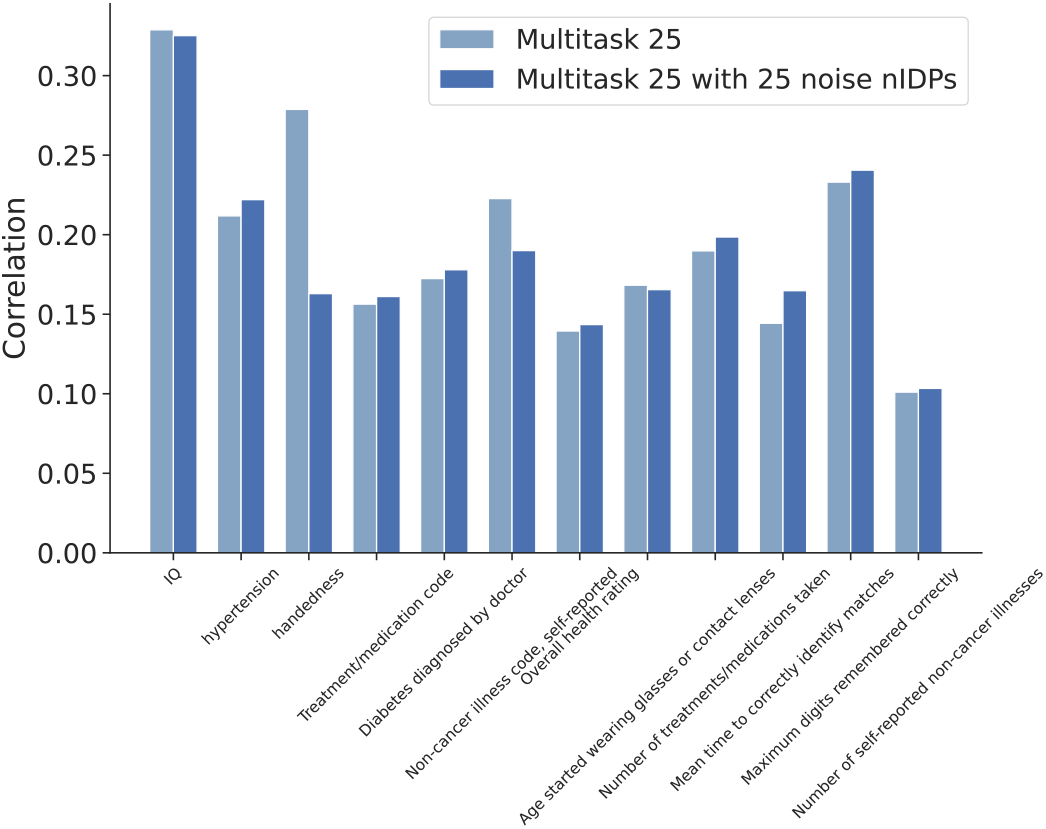
Comparing the prediction performance of multitask SuperBigFLICA with and without uncorrelated nIDPs added in. In the comparison group, we added 25 most uncorrelated auxiliary nIDPs together with 24 most correlated nIDPs for supporting the training of the main nIDP of interests. Results are shown for 12 health outcome and cognitive nIDPs listed in Table 1.

**Fig. 4:**
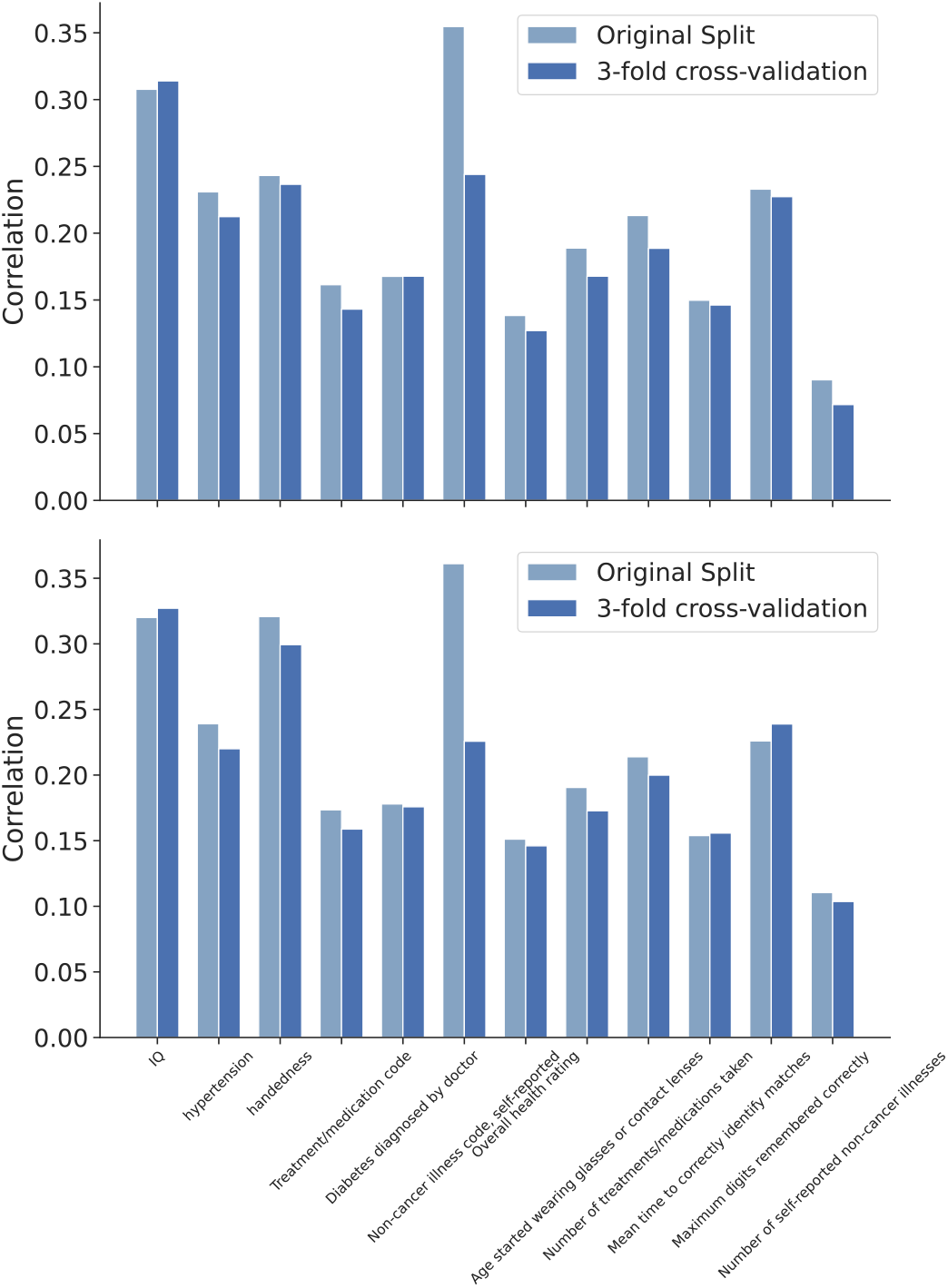
For the 12 health outcome and cognitive nIDPs, comparing the prediction performance of single-task (Top) and multi-task (Bottom) SuperBigFLICA using original fixed train-test split with the combined new settings(1. the dictionary learning preprocessing is performed on the training set; 2. the three-fold cross validation is used.

**Fig. 5:**
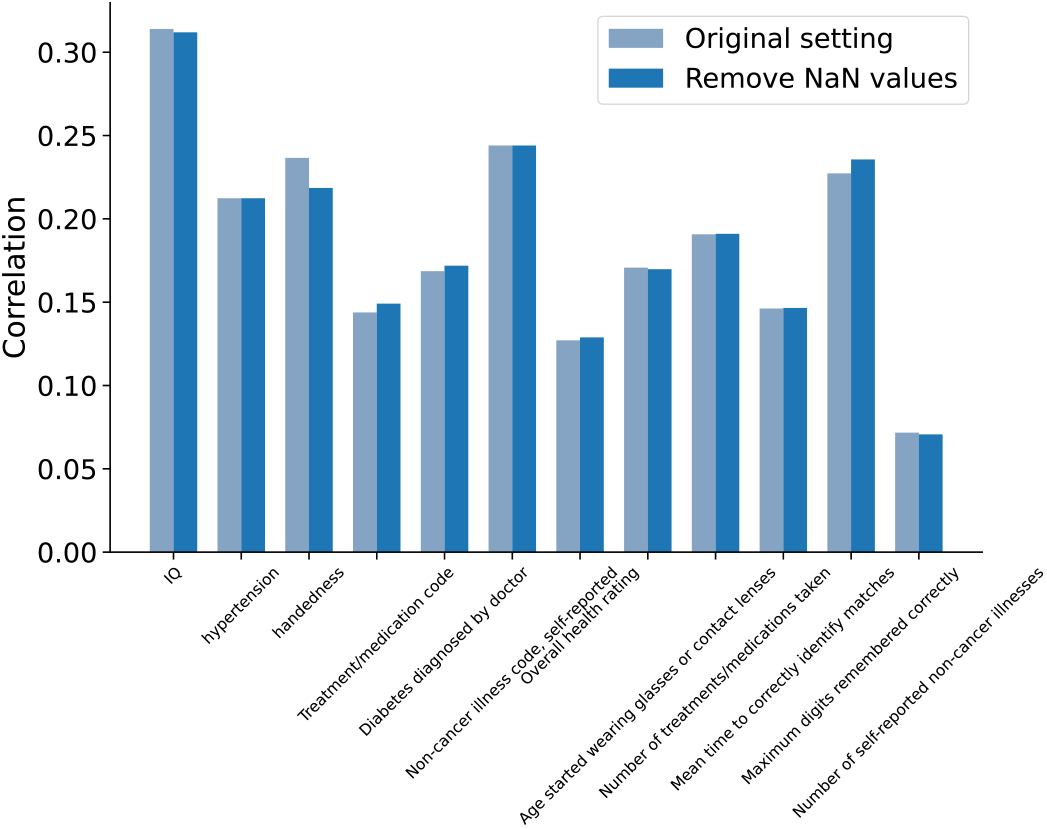
For the 12 health outcome and cognitive nIDPs, comparing the prediction performance of with and without including missing values in the training.

### B. Comparing with existing (semi-)supervised approaches

We compared SuperBigFLICA with two existing (semi-)supervised approaches [13], [16]. **Fig.6** shows the boxplots for the mean correlation of predicting 12 health outcomes and cognitive nIDPs. SuperBigFLICA performs better than MCCAR significantly [13], and almost the same as our implemented versions of Zhou’s approach [16]. Note that our implementation improves the computation efficiency of Zhou’s approach by the mini-batch optimization algorithm. Therefore, we would expect different behaviours from the original implementation.

**Fig. 6:**
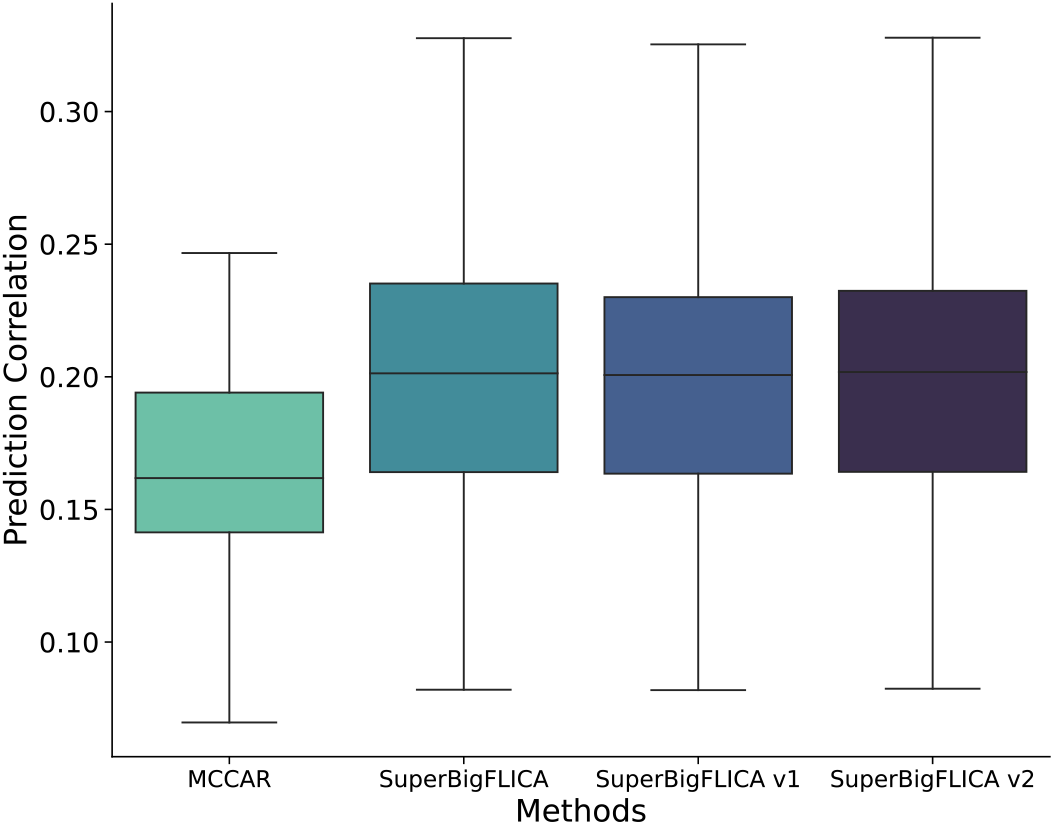
Comparing the mean prediction accuracy of single-task SuperBigFLICA with existing methods on 12 health outcome and cognitive nIDPs listed in Table II. **Boxplot 1:** the prediction accuracy of MCCAR [13]. **Boxplot 2:** the prediction accuracy of SuperBigFLICA. **Boxplot 3:** The prediction accuracy of a variant of SuperBigFLICA, with auto-weighting between different modalities and tasks switched off, i.e., the *σ* ^(*k*)^, *b*^(*k*)^, *γ*_*i*_, *c*_*i*_ parameters are fixed, making the weights equal across modalities and tasks. **Boxplot 4:** The prediction accuracy of another variant of SuperBigFLICA, with both auto-weighting switched off (as box 3) and using an *l*_2,1_-norm regularizer on spatial weights *W* ^(*k*)^ [16].

**Fig.7** futher shows the comparison of prediction correlations of IDPs, BigFLICA, SuperBigFLICA and its variants for all 17,485 nIDPs in UK Biobank (**Fig.7A** and **Fig.7B**). Clearly, BigFLICA outperforms IDPs, and SuperBigFLICA outperforms BigFLICA in many of the nIDP predictions. Our implementation of Zhou’s method [16] based on SuperBigFLICA shows a similar prediction performance as the original SuperBigFLICA, albeit at much increased computational cost (**Fig.7C** and **Fig.7D**).

**Fig. 7:**
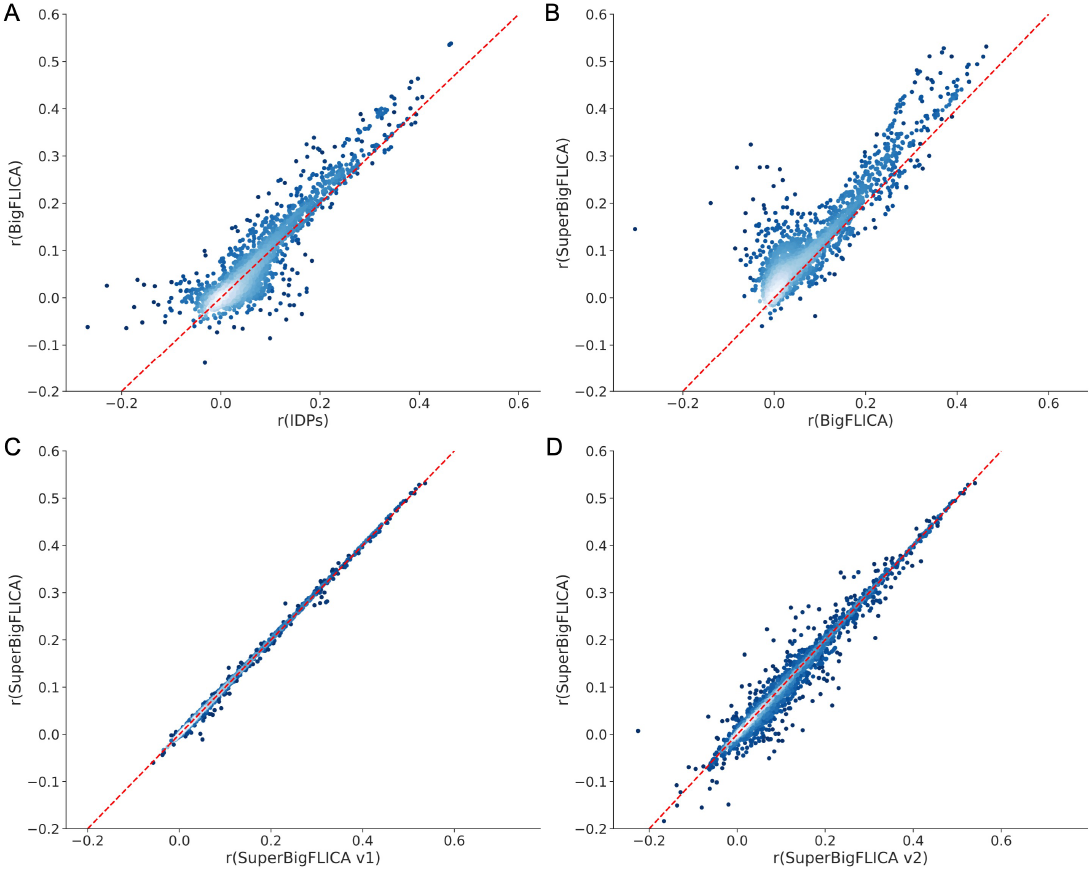
Comparing the prediction correlations using IDPs, BigFLICA, SuperBigFLICA and its variants for all 17,485 nIDPs in the UK Biobank. Comparing the prediction performance of **A:** IDPs vs. BigFLICA; **B:** BigFLICA vs. SuperBigFLICA; **C:** SuperBigFLICA vs. its variants 1 (with auto-weighting between different modalities and tasks switched off); **D:** SuperBigFLICA vs. its variants 2 (with both auto-weighting switched off (as variants 1) and using an *l*_2,1_-norm regularizer on spatial weights *W* ^(*k*)^).

### C. Analysis of SuperBigFLICA algorithm

#### 1) Relationship between prediction accuracy and hyper-parameters

We evaluated the influence of the relative weighting between reconstruction loss and prediction loss, i.e., *λ*, and the number of latent components, on the final prediction performance. The top subfigure of **Fig.8** shows the mean prediction correlation of 12 health outcome and cognitive nIDPs listed in **Table** II across different *λ* and latent components using single-task SuperBigFLICA. When the latent dimension is small, we need to choose a small *λ* (i.e., *λ <* 0.5, which means that the prediction loss has a higher weight than the reconstruction loss) to achieve optimal prediction. When the latent dimension is large, although a smaller *λ* also achieves the best prediction accuracy, the influence of *λ* becomes much smaller than using a smaller latent dimension. We can draw a similar conclusion from the results shown in the middle subfigure of **Fig.8**, which is the case of multi-task SuperBigFLICA with 24 most correlated nIDPs as auxiliary tasks. Conversely, for the generalisability test, the bottom subfigure of **Fig.8** shows that the prediction accuracy is low when the latent dimension is small. When the latent dimension is large, the prediction accuracy is highest when *λ >* 0.5, i.e., the reconstruction loss has a higher weight than the prediction loss. This analysis guides how to select hyper-parameters in different circumstances and demonstrates the usefulness of including both data reconstruction and prediction losses in the objective function.

**Fig. 8:**
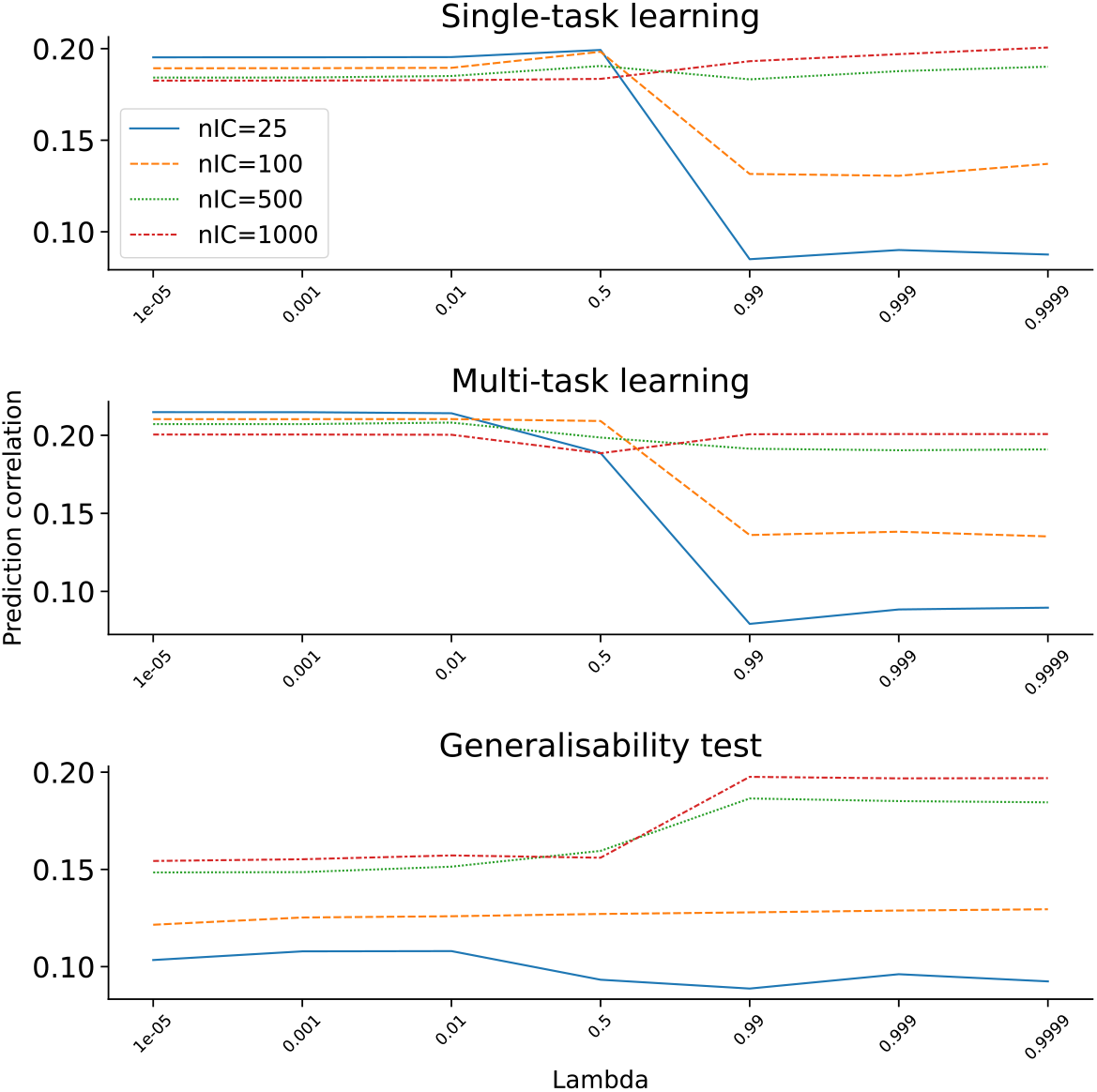
The influence of SuperBigFLICA parameters on the mean prediction accuracy of 12 health outcome and cognitive nIDPs. The relationship between the number of latent dimensions (different line colors) and weights between reconstruction and prediction losses (x-axis) with the mean prediction accuracy across 12 nIDPs listed in **Table** II. **Top**. Single-task learning setting. **Mid**. Multi-task learning setting with 24 auxiliary tasks. **Bot**. generalisability test.

#### 2) Influence of imaging space dimension reduction on prediction accuracy

We first tested whether the imaging space pre-dimension reduction with dictionary learning influenced the final prediction accuracy of nIDPs (Method section III-D). **Fig.9A** shows that, for different nIDPs, the average accuracy of single-task SuperBigFLICA is similar (0.18 *< r <* 0.22) across different dictionary dimensions. The 250-dimensional dictionary learning achieves slightly better performance.

**Fig. 9:**
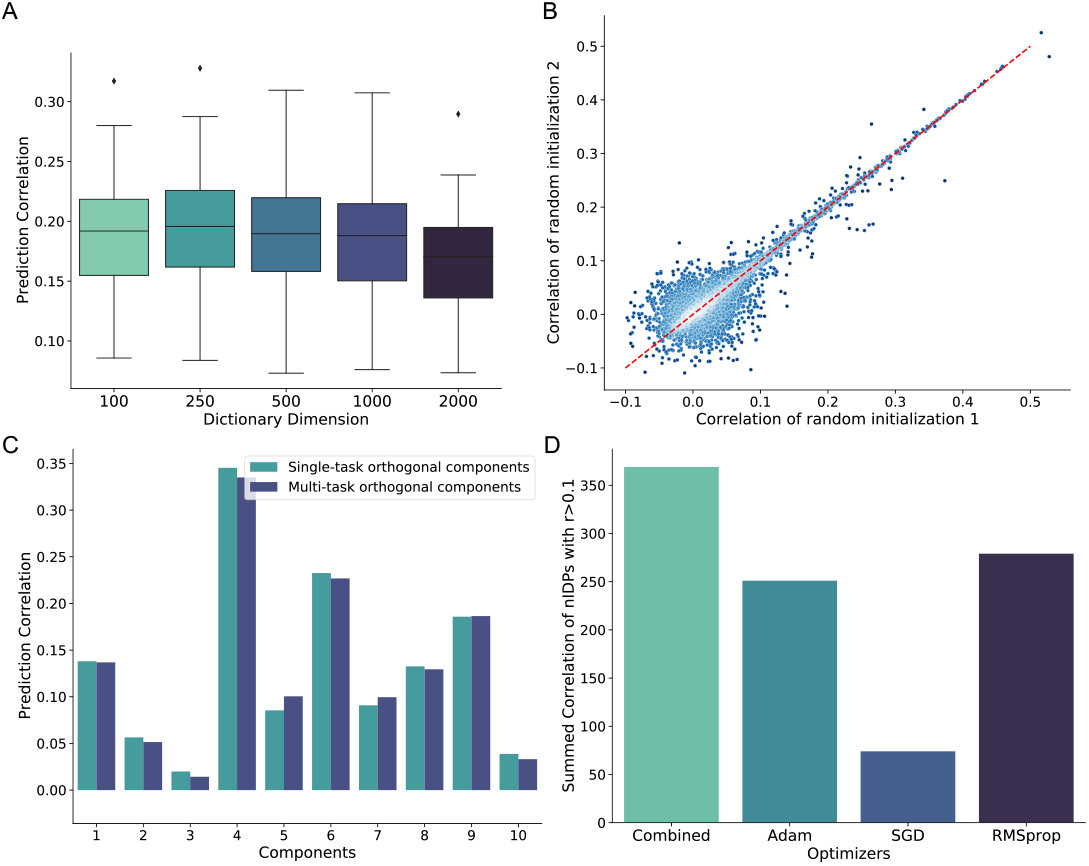
Evaluations of SuperBigFLICA model. **A**. The average prediction accuracy of 12 health outcome and cognitive nIDPs using different dictionary learning dimensions in the image data pre-reduction stage. **B**. The prediction accuracy of 17,485 nIDPs as a result of different random parameter initializations. **C**. Comparing the prediction accuracy of single-task SuperBigFLICA vs. multi-task SuperBigFLICA for predicting top 10 orthogonal principal components derived from all nIDPs. **D**. Comparing the prediction accuracy of all 17,485 nIDPs of SuperBigFLICA optimized by different numerical optimizers.

We further evaluated the use of SuperBigFLICA to perform imaging space dimension “pre-reduction” (250-dimension for each modality, the same with dictionary learning), with all 17,485 nIDPs as supervised targets. We did not see an improvement relative to using dictionary learning, e.g., for fluid intelligence, the best prediction correlation is only around *r =* 0.20, much lower than the result achieved by dictionary learning *r =* 0.32. A lower prediction correlation was also observed for other nIDPs and other SuperBigFLICA dimensions. A possible reason is that using all nIDPs in the pre-dimension reduction stage discards information in the imaging data related to the target nIDP. Also, we find that using SuperBigFLICA in this situation is more memory intensive because we need to keep a huge voxel-by-component spatial weight matrix in memory. In contrast, in dictionary learning, we only need to keep a smaller subject-by-component matrix in memory because it performs a mini-batch optimisation on the voxel dimension. Finally, another disadvantage of the two-stage supervised learning strategy is the need for two nested cross-validation loops for parameter selection, significantly increasing the computation cost.

#### 3) Influence of parameter initialisation on the prediction accuracy

We evaluated the influence of random model parameter initialisations on the final prediction accuracy. We tested whether two different random initialisations will result in different accuracy. Therefore, we performed two multi-task SuperBigFLICA experiments with all 17,485 nIDPs as targets. The only difference between these two experiments is the different seeds for parameter initialisation. **Fig.9B** shows the scatter plots of prediction correlations of the different nIDPs from two different initialisations. We can see those nIDP correlations *<* 0.1 result in a roughly spherical point cloud, while correlations *>* 0.1 are highly correlated, i.e., different initialisations lead to similar nIDP predictions.

#### 4) Influence of nIDP task covariance structure on the prediction accuracy

We further evaluated whether the increases in prediction accuracy of multi-task learning compared with single-task learning result from incorporating information about the task covariance structure into the estimation. We tested this hypothesis by predicting uncorrelated (orthogonal) nIDPs either using separate single-task SuperBigFLICA or jointly using multi-task SuperBigFLICA. To obtain these orthogonal nIDPs, we extracted the top 10 principal components from the subject-by-nIDP data matrix. We compared the accuracy when the model predicted them separately as single-task learning and jointly as multi-task learning. The result shows that the performance is almost the same for each of the 10 “orthogonal” tasks (**Fig.9C**). This experiment shows that the covariance structure between the different nIDPs enables the multi-task model to improve over and above the single-task model. Note that the single-task’s performance is better than the multi-task in some orthogonal principal components. However, the difference is small and not statistically significant, so we believe this is owing to the issue of randomness in the optimizer and parameter initialization.

#### 5) Influence of optimisation algorithms on the prediction accuracy

We finally evaluated the choice of numerical optimisation algorithms on the prediction accuracy. We took an example where we used SuperBigFLICA to fuse all 47 modalities and 17,485 nIDPs to discover a 1,000-dimensional latent space. The Adam, SGD, and RMSprop optimisers all perform worse than the combined optimisation algorithm in terms of the overall prediction accuracy of the 17,485 IDPs (**Fig.9D**). The overall accuracy is estimated by the sum of correlation of nIDPs larger than 0.1. We also tested a full-batch, quasi-newton algorithm L-BFGS but found it always overfits the data, so for briefty, results are not shown here.

#### 6) The execution time of SuperBigFLICA

The time for one iteration of SuperBigFLICA model fitting (training) is between 10s to 30s, depending on whether we predict one nIDP or all nIDPs, using one Intel Skylake 2.6GHZ CPU. SuperBigFLICA usually converges within 50 iterations, so the total running time for analysing the 40k-subject UKB dataset is between 8 mins to 25 mins. Note that we need to use dictionary learning to preprocess the imaging data, which is fast and parallelizable across modalities, taking less than 10 mins per modality for the 40k subjects.

### D. Real data further qualitative analysis

#### 1) Fluid intelligence prediction using a low-dimensional SuperBigFLICA

We first applied a 25-dimensional SuperBigFLICA to predict fluid intelligence scores using data from 47 modalities. **Fig.10A** shows the weights of each of the 25 latent components on predicting fluid intelligence scores (*B* in Eqn. (2)). Components 2, 4, 9, 11, 15, 17, 18, 20, and 23 are selected to contribute to the prediction of fluid intelligence, while the remaining components were switched off by the Laplacian prior to only contributing to the reconstruction of imaging data. The overall prediction correlation is 0.30.

**Fig. 10:**
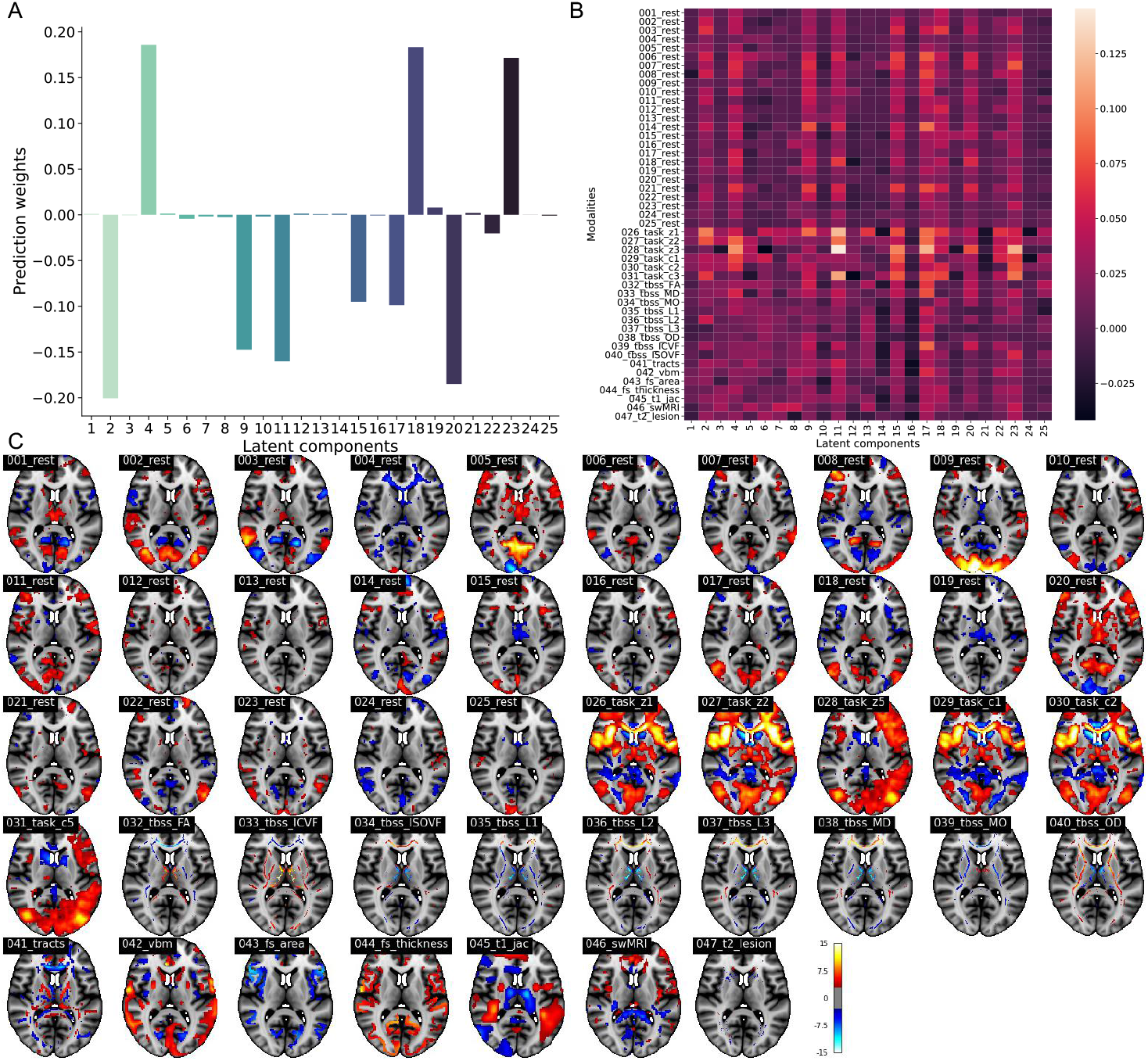
Example results of predicting fluid intelligence with a 25-dimentional SuperBigFLICA. **A**. The prediction weights of fluid intelligence in a 25-dimensional SuperBigFLICA analysis. **B**. The contribution of different modalities within each of 25 SuperBigFLICA components for predicting fluid intelligence. **C**. The *Z*-score normalized 47 spatial maps of modes with strongest contribution to the prediction of fluid intelligence in a 25-dimensional SuperBigFLICA decomposition, with fluid intelligence as the supervision target (component 2, MNI152 coordinate z=10).

**Fig.10B** shows the contribution of each component and each modality to the prediction of fluid intelligence (Method section II-C)). The components selected for fluid intelligence prediction have higher overall contributions, which demonstrated the validity of the proposed way to estimate modality contribution. Across the different modalities, the components from task contrast maps have the highest contribution to the prediction of fluid intelligence, while the resting-state dual-regression spatial maps have a slightly lower contribution. The dMRI and sMRI derived modalities contribute the least to the prediction.

**Fig. 10C** shows the *z*-score spatial maps of component 2 for each modality, which was generated by regressing latent components back onto the original voxel-wise data. This component contributes most to the prediction of fluid intelligence scores. The task contrast maps and resting-state map 5 have the highest voxel-wise activations. The regions that show the highest contribution to the prediction of fluid intelligence are mainly located in the precuneus cortex, posterior cingulate gyrus, lateral occipital cortex, insular cortex, inferior frontal gyrus, and frontal pole [54]. Among them, the insular cortex, inferior frontal gyrus, and frontal pole were found significant in task modalities in our previous BigFLICA approach [2], but with a much higher 750-dimensional decomposition. This reflects the fact that adding supervision to the model can help the model learn task-specific patterns easier. In addition, here, with the SuperBigFLICA approach, we can observe a more comprehensive ‘multimodal’ effect, such as the changes in cortical surface area and thickness in these regions, and changes in tracts and white matter microstructures that connect these regions, which are also reported in literature [55], [56]. Moreover, we also observe the precuneus cortex and posterior cingulate gyrus in several resting-state maps, as part of the default mode network, involved in fluid intelligence prediction [57].

#### 2) Phenotype discovery using a high-dimensional SuperBigFLICA

We finally applied SuperBigFLICA to perform a 1,000-dimensional decomposition with all 17,485 nIDPs as supervision targets. This 1,000-dimensional latent space can serve as a set of new data-driven IDPs.

**Fig. 11** shows the *Z*-score normalised spatial maps of the component that most strongly contributes to the prediction of *hypertension* scores. The prediction correlation is 0.24. The regions that show the highest contribution to the prediction of hypertension are mainly located in the precuneus cortex, visual cortex, middle temporal gyrus, central opercular cortex, Heschel’s gyrus, inferior frontal gyrus and insular cortex, and also external capsule tracts. Again, the modes are more ‘multimodal’, and several consistent findings have been reported in the literature [58]–[60].

**Fig. 11:**
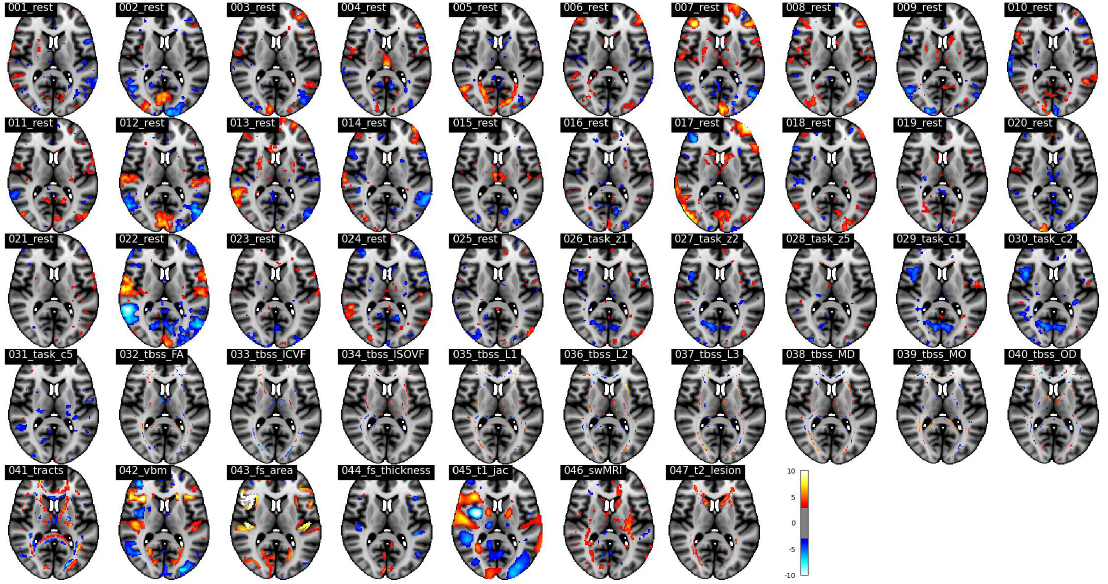
The *Z*-score normalized spatial maps of the strongest modes that contributing to the prediction of *hypertension* in a 1,000-dimensional SuperBigFLICA, with all 17,485 nIDPs as supervision targets (MNI152 coordinate z=10).

Likewise, **Fig. 12** shows the *Z*-score normalised spatial maps of the mode that contributes most strongly to the prediction of *age started wearing glasses or contact lenses*. The prediction correlation is 0.19. The regions that show the highest contribution to the prediction of hypertension are mainly located in visual areas, especially for resting-state dual regression spatial maps 5, which represents the visual network.

**Fig. 12:**
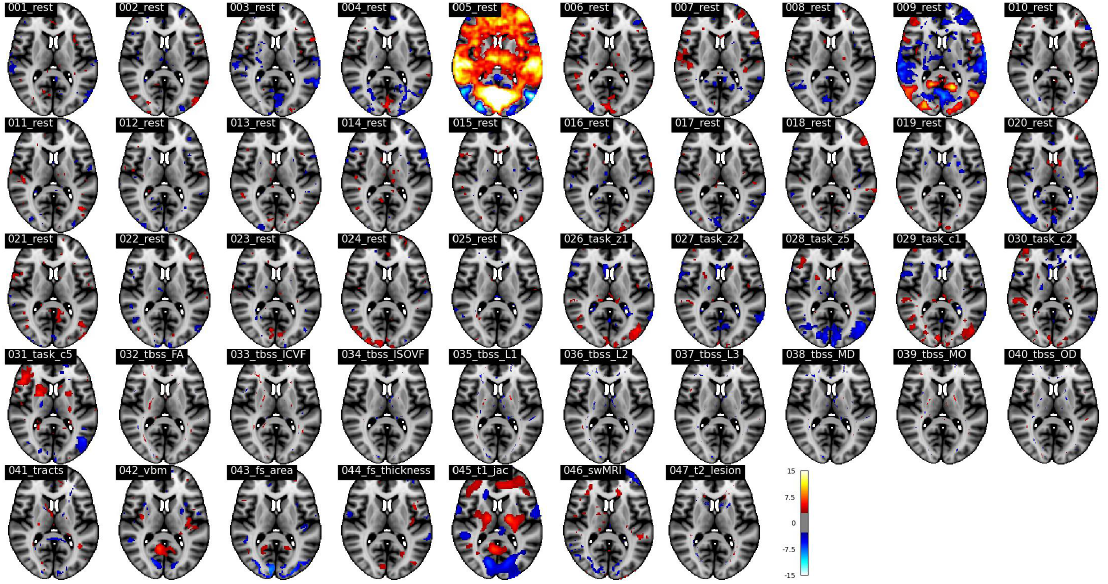
The *Z*-score normalized spatial maps of the strongest modes that contributing to the prediction of *age started wearing glasses or contact lenses* in a 1,000-dimensional SuperBigFLICA, with all 17,485 nIDPs as supervision targets (MNI152 coordinate z=10).

## V. Discussion

In this paper, we propose SuperBigFLICA, a semi-supervised multimodal data fusion approach that simultaneously reconstructs the original voxel-wise imaging data and best predicts non-imaging derived phenotypes. The approach is scalable to extreme high-dimensional data sets, e.g., UK Biobank scale neuroimaging datasets. SuperBigFLICA inherits the Bayesian framework from the previous FLICA model [2], [8]. Additionally, it incorporates an additional prediction term to enable supervised learning of the target variable of interests (i.e., multiple nIDPs). SuperBigFLICA can discover spatially sparse and orthogonal modes that can serve as generic data-driven IDPs for future prediction of new nIDPs. The weighting of different modalities and nIDPs can be automatically inferred from the data, avoiding manual specification.

Compared to previous linear approaches (e.g., [13]), the scalability of our approach to huge data sets is improved through the use of advanced stochastic optimisation algorithms. Our model can use multiple nIDPs as supervision targets and can predict unseen nIDPs. Compared to nonlinear approaches (e.g., [16]–[18]), our approach can explicitly discover a low-dimensional linear latent space as new image-derived phenotypes. We performed a comprehensive comparison of SuperBigFLICA with the hand-curated IDPs currently being created by our group on behalf of UK Biobank, and modes of unsupervised BigFLICA, and found a significantly improved performance on predicting nIDPs. We also showed that by using the multi-task learning paradigm, SuperBigFLICA showed a further improvement than its single-task setting. We demonstrated SuperBigFLICA’s performance in learning a generalisable latent space by applying it to predict unseen nIDPs. These tests were performed using the largest neuroimaging dataset to date (UK Biobank), with 47 different modalities, 39,770 subjects, and 17,485 nIDPs, which illustrates the ability of SuperBigFLICA for analysing large-scale datasets. In real data examples, we demonstrated that SuperBigFLICA finds interpretable modes predictive of health outcomes and cognitive nIDPs.

There are multiple future directions for improving the current approach. First, we could further explore the possibility of improving the prediction of unseen nIDPs by using advanced techniques in transfer learning [61]. Second, a deeper understanding of the latent space, including the interpretation of spatial maps and the influence of dimensionality of latent space with prediction power, could be interesting. Third, another straightforward extension would be adding nonlinearity to SuperBigFLICA, which enables it to extract more complex nonlinear patterns from brain imaging data, with or without the supervision of nIDPs. Many options exist to achieve this by using either deep neural networks [62] or traditional machine learning approaches such as Gaussian process latent variable model [63] and multiple kernel learning [64]. Nonlinear approaches such as deep convolutional neural networks have shown excellent age and sex prediction accuracy using structural MRI data and in Alzheimer’s disease progression [65], but the usefulness of nonlinear models for neuroimaging data is still under debate [66]–[68] due to the increased complexity of evaluating and interpreting their performance. Therefore, besides developing a nonlinear model for improving predictive performance, deriving interpretable nonlinear features is also an important task.

In summary, all of the above will be explored in future improvements to our analysis approach. An easy-to-use version of this software will be integrated into an upcoming version of the FSL software package [45], [46]. Results from applying SuperBigFLICA on UK Biobank will also be released via the UKB database as new data-driven IDPs (image features), further contributing to the richness of the sample and enabling neuroscientific research.

## Notes

Financial support was provided by the Wellcome Trust Core Award Grant Number 203141/Z/16/Z, the Wellcome Trust Collaborative Award in Science 215573/Z/19/Z and further supported by the Netherlands Organization for Scientific Research Vici Grant No. 17854 and NWO-CAS Grant No. 012-200-013. We are grateful to UK Biobank and its participants (access application 8107). Computation used the Oxford Biomedical Research Computing (BMRC) facility, a joint development between the Wellcome Centre for Human Genetics and the Big Data Institute supported by Health Data Research UK and the NIHR Oxford Biomedical Research Centre. The authors declare that they have no competing financial interests.

### Competing Interest Statement

The authors have declared no competing interest.

### Summary of Updates

1

